# Lamellar junctions in the endolymphatic sac act as a relief valve to regulate inner ear pressure

**DOI:** 10.1101/143826

**Authors:** Ian A. Swinburne, Kishore R. Mosaliganti, Srigokul Upadhyayula, Tsung-Li Liu, David G. C. Hildebrand, Tony Y.-C. Tsai, Anzhi Chen, Ebaa Al-Obeidi, Anna K. Fass, Samir Malhotra, Florian Engert, Jeff W. Lichtman, Tom Kirchhausen, Eric Betzig, Sean G. Megason

## Abstract

The inner ear is a fluid-filled closed-epithelial structure whose normal function requires maintenance of an internal hydrostatic pressure and fluid composition by unknown mechanisms. The endolymphatic sac (ES) is a dead-end epithelial tube connected to the inner ear. ES defects can cause distended ear tissue, a pathology often seen in hearing and balance disorders. Using live imaging of zebrafish larvae, we reveal that the ES undergoes cycles of slow pressure-driven inflation followed by rapid deflation every 1-3 hours. Using serial-section electron microscopy and adaptive optics lattice light-sheet microscopy, we find a pressure relief valve in the ES comprised of thin overlapping basal lamellae that dynamically extend over neighboring cells before rupturing under pressure leading to ES collapse. The unexpected discovery of a physical relief valve in the ear emphasizes the need for further study into how organs control fluid pressure, volume, flow, and ion homeostasis in development and disease.

## Introduction

Understanding the mechanisms by which organs use water-filled cavities to compartmentalize biochemical and biophysical environments is a fundamental problem. Because water is nearly incompressible, several organs harness and cope with water as an object that transmits force. Hydrostatic pressure inflates the eye during development (Coulombre, 1956). Later, unstable ocular pressure from reduced production of aqueous humor or reduced drainage can lead to blindness, as occurs in hypertonic maculopathy or glaucoma (Costa & Arcieri, 2007; Leske, 1983). Hydrostatic pressure appears to drive expansion of brain ventricles during development (Desmond, 1985; Desmond & Levitan, 2002; Lowery & Sive, 2005). Later, unstable hydrostatic pressure in brain ventricles is correlated with hydrocephaly and mental disorders (Hardan, Minshew, Mallikarjuhn, & Keshavan, 2001; Kurokawa et al., 2000). We have recently shown that hydrostatic pressure inflates and controls the size of the developing zebrafish ear (Mosaliganti et al., under review). Later, unstable pressure in the ear can cause deafness and balance disorders like Pendred syndrome and Meniere’s disease (Belal & Antunez, 1980; Schuknecht & Gulya, 1983). This theme of harnessing hydrostatic pressure for normal development and controlling pressure for healthy physiology raises the question of how tissues regulate pressure. Tissue structures identified as important for pressure control include Schlemm’s canal in the eye, arachnoid granules and the choroid plexus in the brain, and the endolymphatic duct and sac in the ear. A cavity’s pressure could be managed via mechanisms involving molecular pores, molecular transporters, and the physical behavior of the tissue. Observing the mechanisms by which these tissue barriers control pressure has been limited by a range of obstacles such as optical accessibility as well as uncertainty regarding both the time- and length-scale on which they function. As such, not much is known about how these tissues manage fluctuating pressures because they have not been observed in vivo.

The inner ear is a prominent example of an organ whose tissue form determines its physiology. The inner ear is filled with endolymph whose composition differs from other fluids such as plasma, perilymph, and cerebral spinal fluid with its high potassium, low sodium, and high electric potential (Lang, Vallon, Knipper, & Wangemann, 2007). This endolymph composition is necessary to drive ion currents in hair cells to convert fluid movement, driven by either the head’s acceleration or by sound, into a biochemical signal. The ear’s endolymphatic duct connects the endolymph in the semicircular canals, cochlea, and other chambers to the endolymphatic sac (ES, Figure 1A,B, blue square highlight in B). An ES-like structure is present in basal vertebrates, including lamprey and hagfish (Hammond & Whitfield, 2006), suggesting an ancient role in inner ear function. The epithelium of the ES, as well as of the rest of the ear, has an apical surface facing the internal endolymph and a basal surface facing the external perilymph (beige endolymph, magenta perilymph, Figure 1A). Excessive hydrostatic pressure can tear the epithelium, disrupting the electric potential (a pathology called endolymphatic hydrops). In contrast, low potassium or reduced endolymph production can lower hydrostatic pressure within the ear to the point where the ear chambers may collapse (Lang et al., 2007). Early work suggests that the ES has an important role in endolymph homeostasis: if the ES is ablated, then hydrostatic pressure rises and the epithelium can tear (Kimura, 1965; Naito, 1950). While many molecular pores and transporters are expressed in the ES, it is unclear why it is organized as a deadend tube.

**Figure 1.**
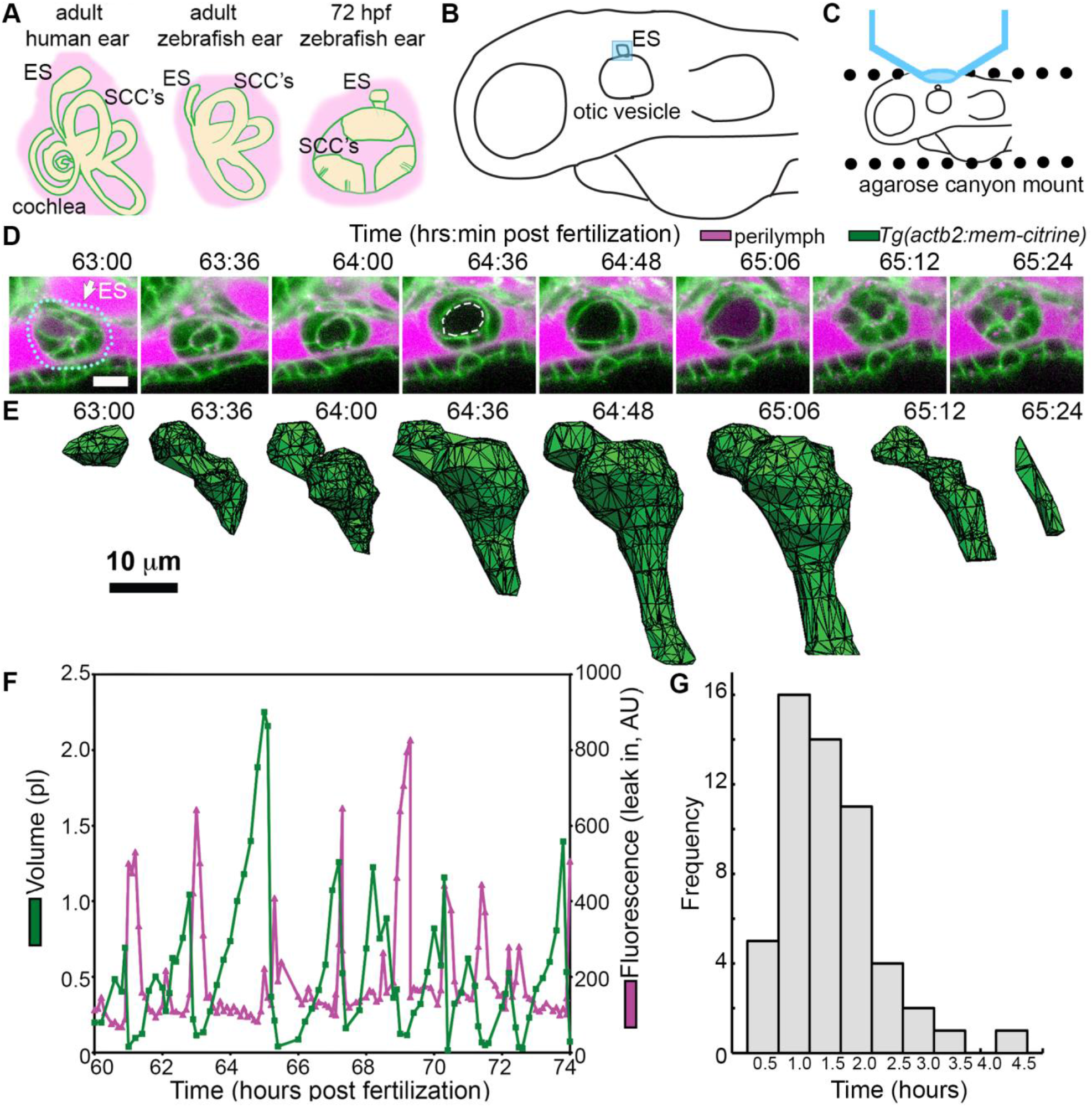
ES lumen slowly inflates and rapidly deflates every 1–3 hours. (**A**) Illustration of the adult human inner ear showing cochlea, semicircular canals (SCC’s), and endolymphatic duct and sac (ES). Illustrations of the adult and larval zebrafish inner ear highlight position of ES relative to otic vesicle, perilymph (magenta), and endolymph (beige, see also Figure—figure supplement 1 and Video 1 for how the zebrafish ES first forms). (**B**) Illustration of larval zebrafish that presents context of ear (lateral perspective, ES and microscopy region highlighted with blue square). (**C**) Illustration of imaging setup. (**D**) Slices and select time points from 3D confocal time course showing a single inflation and deflation event from a live zebrafish embryo. Cell membranes (green) are labeled using ubiquitous membrane citrine transgenes. Perilymph (magenta) is labeled with 3 kDa dextran-Texas red. ES identified with dotted blue outline and white arrow. Lumen of inflated ES identified with dashed white outline in 64:36 panel. (**E**) Corresponding 3D meshes of the segmented ES lumen volume. (**F**) Quantification of segmented ES volumes (primary axis, green) and leak in fluorescence (secondary axis, magenta) over multiple cycles (see also Figure 1—figure supplement 1B,C and Videos 2–3). (**G**) Histogram of times between peak inflation volumes, compiled from 8 different time courses and 54 inflations. Scale bars 10 μm.

The ES has not been studied in zebrafish because it has not always been clear that zebrafish have an ES or, that if zebrafish have one, at what time it forms during development (Haddon & Lewis, 1996; Hammond & Whitfield, 2006). In situ hybridization images at 52 hours post fertilization (hpf) showing expression of ES markers, *foxi1* and *bmp4*, restricted to an ES like structure are the strongest and developmentally earliest evidence for zebrafish having an ES (Abbas & Whitfield, 2009; Geng et al., 2013), but its formation and physiological function remained unknown. Here we present a detailed characterization of the ES in zebrafish performed using confocal microscopy, serial-section electron microscopy, adaptive optics lattice light-sheet microscopy, and a genetic mutant that together demonstrate that the ES contains a physical relief valve for regulating inner ear pressure.

## Results

### The endolymphatic sac exhibits cycles of inflation and deflation

We first established an imaging system to identify the developmental origins of the zebrafish ES. Extended time lapse imaging required long-term immobilization with α-bungarotoxin that permits normal development without the reduction in ear growth caused by long-term treatment with tricaine, bright fluorescent transgenic fish with contrast from a membrane localized fluorescent protein, and mounting in a submerged agarose canyon that permits positioning of the zebrafish ear close to the coverslip (400 μm wide, walls of canyon secure the yolk and head, Figure 1C) (Swinburne, Mosaliganti, Green, & Megason, 2015). The zebrafish ear develops from a collection of epithelial cells that form a closed fluid-filled ellipsoidal structure called the otic vesicle (Figure 1B) (Whitfield, 2015). We saw that ES morphogenesis begins at 36 hpf as an evagination in the dorsal epithelial wall of the otic vesicle (Figure 1—figure supplement 1A, Video 1, dorsal region or interest highlighted with blue box, Figure 1B). Between 36-60 hpf, the ES grows, elongates, and moves into a more central and medial position due to morphogenesis of the otic vesicle epithelium. This position is consistent with the prior in situ patterns of ES markers. These findings established that the zebrafish ES is optically accessible during embryonic and larval stages and that ES morphogenesis begins at 36 hpf.

The zebrafish ear becomes physiologically functional between 60 and 72 hpf, as demonstrated by the onset of the vestibulo-ocular reflex (Mo, Chen, Nechiporuk, & Nicolson, 2010). We found that the ES begins to exhibit a physical behavior during the same time window. We observed that the lumen of the ES remains closed until 60 hpf, but between 60 and 65 hpf it begins cycles of slowly inflating and rapidly deflating (Figures 1D, E, Figure 1—figure supplement 1, and Videos 2–3, yellow arrow points to ES, blue dotted line traces basal surface of ES, white dashed line traces apical interface with ES lumen, sagittal slices correspond to region highlighted with blue box in 1B). Three-dimensional (3D) measurements showed that the ES lumen volume changes 5–20-fold through the course of each cycle (Figure 1E, F, volume plotted in green, and Figure 1—figure supplement 1B-C). The period between peak ES volumes exhibited a broad distribution (0.34–4.45 hours) with an average of 1.59±0.77 hours (mean±SD, histogram compiled from 8 time courses and 54 peaks, Figure 1G). These observations allow for several potential causes of the inflation-deflation cycles of the ES: a response to organ-wide increases in fluid pressure within the otic vesicle; a local tissue behavior where the ES inflates with fluid from the perilymph; or a local tissue behavior where cells in the ES periodically coordinate their movements to expand the ES volume.

### Loss of the epithelial barrier is sufficient for ES deflation

To assess whether ES inflations occur by transmission of endolymph pressure between the otic vesicle and the ES, we injected a small volume of solution containing 3 kDa fluorescent dextran into the otic vesicle and followed its movement during inflation and deflation. We found that the duct is open and that endolymph flows from the otic vesicle into the ES during inflation (Figure 2A-F, Video 4). Additionally, upon deflation, endolymph rapidly leaked out of the ES and into the perilymphatic space (75:22 panel, Figure 2E, F). To determine whether pressure in the otic vesicle is transmitted to the ES for its inflation, we laser-ablated 2-3 cells within the wall of the otic vesicle distant from the ES at 64 hpf (Figure 2G). Shortly after this treatment, the ES deflated and completely collapsed within 20 minutes (Figure 2H). Together, these results indicate that: (1) fluid pressure in the otic vesicle inflates the ES volume; (2) the ES tissue has elastic material properties; and (3) a loss of epithelial integrity is sufficient for ES deflation.

**Figure 2.**
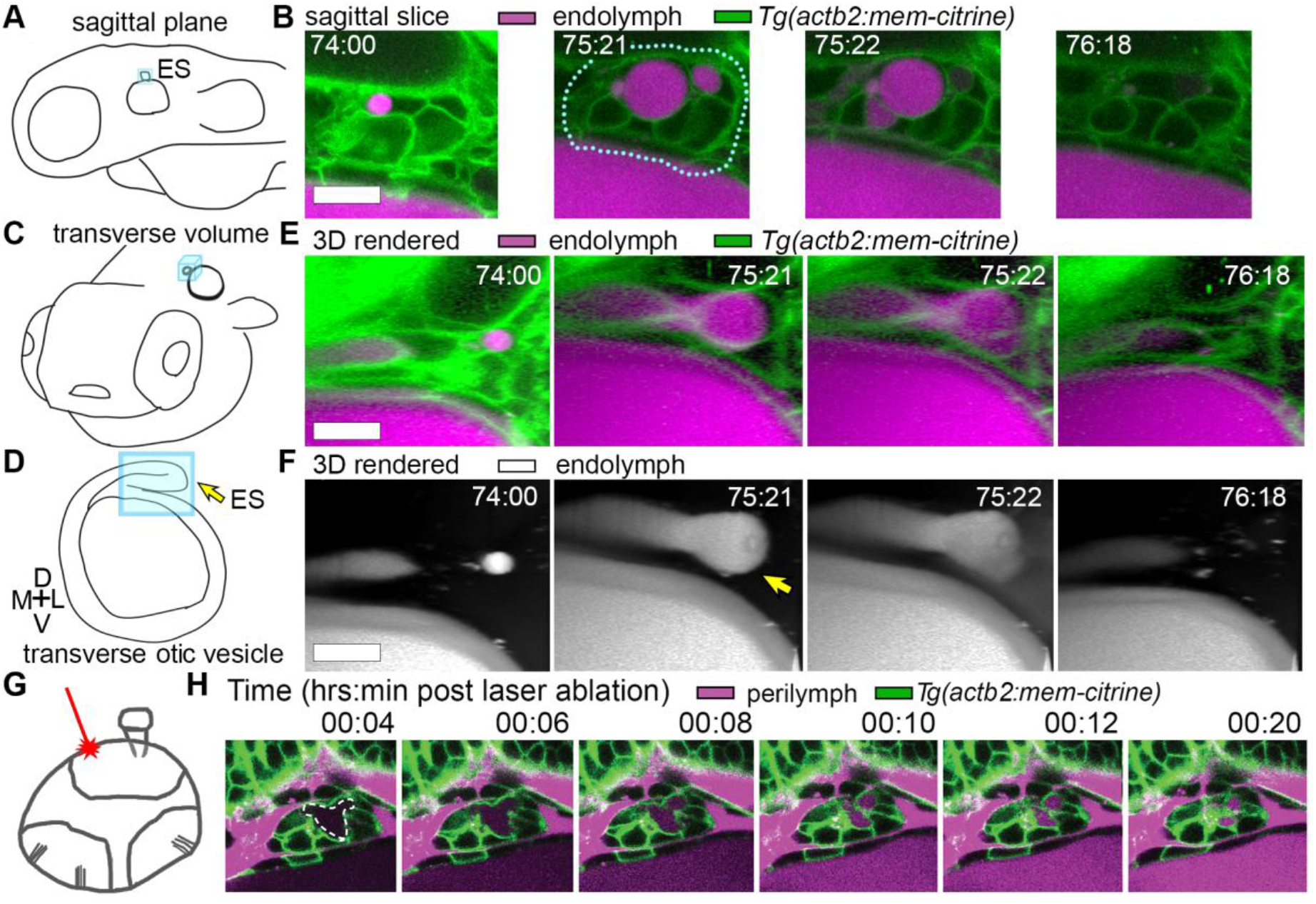
Hydrostatic pressure transmits endolymph through duct to inflate the ES. (**A**) Illustration of larval zebrafish highlighting sagittal plane of image acquisition (blue square). (**B**) Time points of individual sagittal slices of raw data from 3D time course (endolymph labeled magenta by dye injection, membrane citrine in green). ES identified with dotted blue outline. (**C**) Illustration of larval zebrafish highlighting transverse perspective (blue box) for rendered volumes of ES. (**D**) Illustration of otic vesicle highlighting a transverse plane of duct exiting dorsal medial wall of vesicle, ending in ES. Blue square highlights regions rendered in E and F. Yellow arrow points to ES. (**E**) Time points from time course of raw data rendered in 3D transverse view (endolymph in magenta, membrane-citrine in green) showing endolymph flowing through duct to ES and then out to perilymph (see also Video 4). (**F**) Same renderings with only endolymph in grey. Yellow arrow points to ES lumen inflated with endolymph. (**G**) Illustration of strategy for laser ablating otic vesicle cells with point scanning 2-photon laser to ablate 2-3 cells at target. (H) Slices from 4D confocal time course after laser ablation showing ES deflation. ES lumen is outlined with a white dashed line in (**H**). All scale bars 10 μm.

A core function of epithelial sheets is to act as a barrier that can prevent passive movement of molecules between an organ’s interior and exterior. Deflations could be driven by local breaks in the epithelial barrier or by an alternative mechanism such as cellular reabsorption of endolymph. To distinguish these mechanisms, we used dye injections to label the perilymph that surrounds the otic vesicle and ES (magenta, Figure 1D). Quantification of fluorescence within the lumen of the ES reveals that perilymph dye begins to leak into the ES lumen at the onset of each deflation (magenta plot, Figure 1F, visible leakage in Figure 1D, Videos 2-3). In the example shown, for each of 9 major rounds of inflation and deflation, deflation coincided with an influx of fluorescence from the perilymph into the lumen of the ES. This coincidence of timing suggests that both deflation and dye leakage were due to transient breaks in the epithelial barrier. This interpretation is consistent with the rapid release of endolymph prior to deflation (Figure 2E, F, Video 4). However, because of the limit of our spatial resolution as well as the absence of contrast for trafficking vesicles, we could not dismiss alternative mechanisms such as large amounts of rapid transcytosis.

The observations that perilymph enters the deflating ES may seem counterintuitive since the contents of a punctured high-pressure elastic vessel would be expected to primarily flow outwards. To determine whether diffusion could explain the “upstream” movement of dye from the perilymph to the ES, we estimated the Péclet number, which is the ratio of advective transport rate (originating from bulk flow) to diffusive transport rate (originating from the random walk of Brownian motion). A large Péclet number indicates that bulk flow dominates transport while a low Péclet indicates a significant contribution from diffusive movement. We analyzed the deflation kinetics, to estimate an average advective flow velocity, originating from bulk movement created by the elastic collapse of the ES, of ~0.028 μm/s. To estimate the contribution of diffusive transport we assumed a 5 μm flow length, based on measuring the thickness of cell bodies in the ES, and a dye diffusion coefficient of ~120 μm^2^/s. The calculated ratio, or Péclet number, of 0.001 indicates that diffusive movement dominates over advection, thus explaining the observed flux of dye from perilymph into the ES and supporting the interpretation that it represents a transient break in the epithelial barrier. It is difficult to explain the simultaneous inward and outward movement of fluid with the alternative interpretation that rapid transcytosis deflates the ES.

### Lmx1bb is essential for ES deflation

The uniqueness of the ES inflation and deflation cycles suggests specific genes might contribute to the development of this physiology. To identify pathways that contribute to the emergence of this physiology and to reveal ways in which it can malfunction, we examined a mutant with an enlarged ES. Previously, we mapped a zebrafish ear mutant as being caused by a premature stop codon in the transcription factor *lmx1bb* (Obholzer et al., 2012). In zebrafish *lmx1bb* mutants the ES became greatly enlarged (greater than four times wild-type volume) making it readily visible at 80 hpf by bright-field microscopy (Figure 3A, white asterisk). To determine if *lmx1bb* is expressed at an appropriate place and time for a mutation to be causing an ES defect, we imaged a transgenic reporter line driven by the *lmx1bb* promoter—*Tg(lmx1bb:eGFP)^mw10^*. This reporter was expressed in ES cells beginning at 52–58 hpf, just before the first ES inflation (Figure 3B), and thus consistent with *lmx1bb* expression being instrumental for development of the ES’s ability to release pressure. Live imaging and perilymph tracking in *lmx1bb^jj410/jj410^* mutant embryos revealed that the ES diffusion barrier does not break and the lumen continually inflates (Figures 3C-D, Figure 3—figure supplement 1, and Videos 5–6). Together, these results indicate that *lmx1bb* is necessary for breaks in the ES’s epithelial barrier and that absence of normal ES deflation physiology can lead to a state like endolymphatic hydrops.

**Figure 3.**
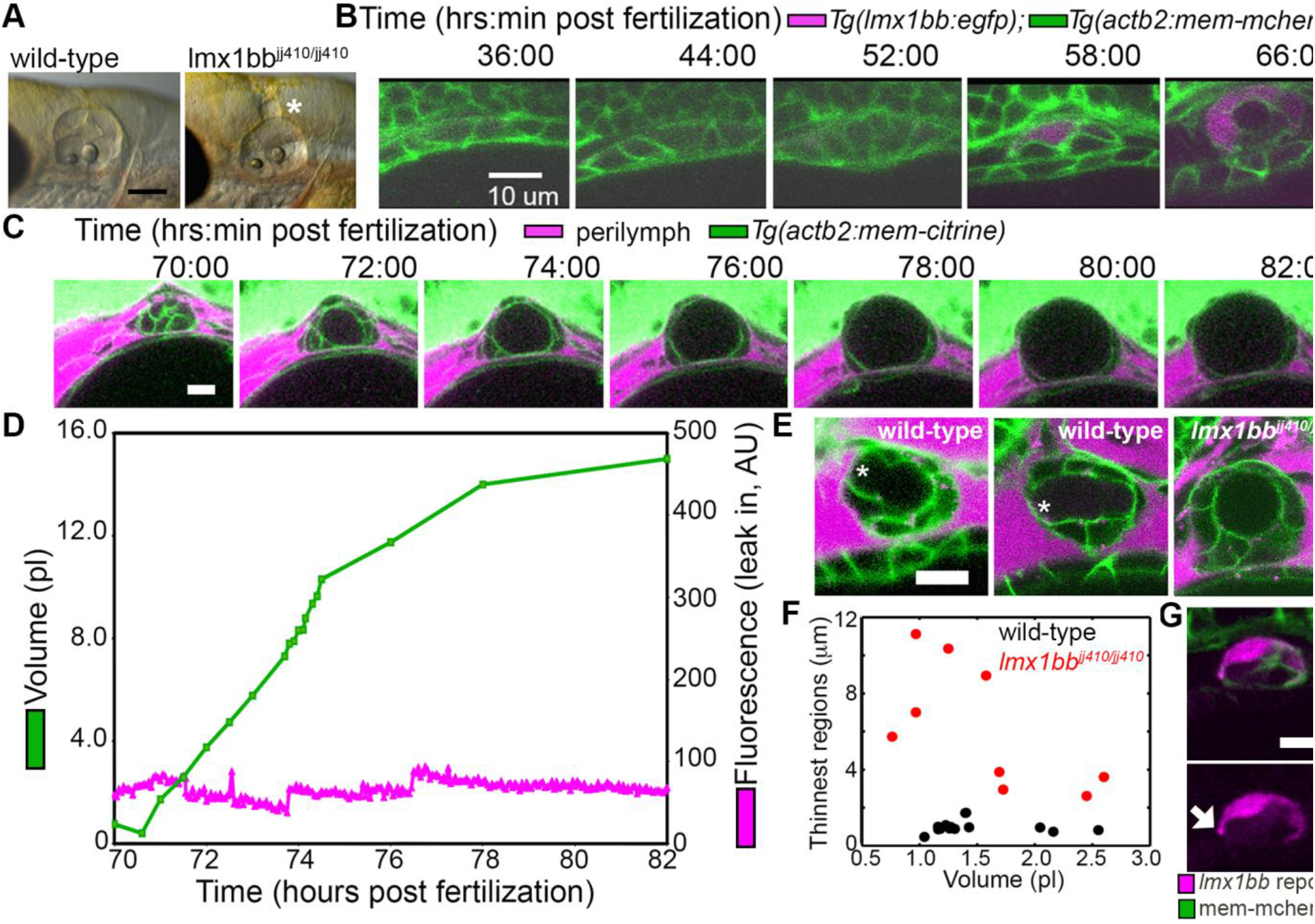
Lmx1bb is necessary for development of the ES’s ability to form breaks in its diffusion barrier and deflate. (**A**) Lateral view of wild-type and *lmx1btì^jj410/jj410^* mutant ears imaged by bright-field microscopy at 80 hpf, asterisk labels greatly enlarged mutant ES. Scale bar 100 μm. (**B**) Slices from 3D confocal time course of an *lmx1bb* transcriptional reporter (magenta, *Tg(lmx1bb:egfp)^mw10/mw10^*, green, *Tg(actb2:mem-mcherry2)^hm29^*. (**C**) Slices and select time points from 3D confocal time course of *lmx1bb^jj410/jj410^* mutant embryos. Membrane (green) from ubiquitous membrane citrine transgenes. Perilymph (magenta) from 3 kDa dextran-Texas red. (**D**) Quantification of segmented ES volumes (primary axis, green) and leak in fluorescence (secondary axis, magenta) from *lmx1bb^jj410/jj410^* time course in C (see also Figure 3—figure supplement 1 and Videos 5–6). (**E**) Small regions with thin membranes (asterisks) form in the inflated ES of wild-type but not *lmx1bb* mutants. (**F**) Quantification of minimum epithelial thickness versus inflated ES volume in mutant (plotted in red, N=9) and wild-type (plotted in black, N=14). Compiled from embryos between 65-80hpf. (**G**) Mosaic labeling from *Tg(lmx1bb:egfp)* reveals thin basal processes, indicated with white arrow. All scale bars in (**B-G**) are 10 μm.

### The ES relief valve is made of basal lamellar junctions

The absence of ES deflation in the *Imx1bb* mutant suggested a structural deficiency. Comparing the dorsal ES tissue between wild-type and *Imx1bb* mutants using confocal microscopy and membrane localized fluorescent proteins, we observed that the ES tissue was much thinner in wild-type embryos during peak inflation than in the corresponding region in mutants inflated to a similar volume (thin region indicated by asterisk, Figure 3E, F, data in F compiled from inflation events occurring between 65 and 80 hpf). In wild-type embryos this region often appeared as thin as a single membrane (1.0±0.3 μm, mean ± SD) rather than being thick enough to identify the characteristic apical and basal sides of an epithelium, which were still present in *Imx1bb* mutants (6.2±3.2 μm, the mutant, N=9, is significantly thicker than wild-type, N=14, with a Mann-Whitney-Wilcoxon one tailed p-value of 4 × 10^−5^). Furthermore, imaging the mosaic signal of the *Tg(Imx1bb:eGFP)* reporter revealed thin basal protrusions that extend along the basal side of neighboring ES cells (white arrow, Figure 3G). The absence of local breaks in the diffusion barrier in combination with the absence of regional thinning of cells in the inflated ES of *lmx1bb* mutants suggested that these thin areas might be structurally relevant to pressure relief in the ES.

We determined the ultrastructure of these thin areas using serial-section electron microscopy (EM) at a resolution of 4.0 × 4.0 × 60 nm^3^ per voxel. We found the dorsal ES tissue of the inflated wild-type sac is an extremely thin shell of partially overlapping lamellae extending from the basal sides of multiple adjacent cells (Figure 4; Video 7). The basal extensions appear to be the distal barrier in the ES because the endolymph they contain is continuous with the lumen of the ES and endolymphatic duct (lumen extends ventrally from outlined ES lumen in third panel of Figure 4A). We term these “lamellar junctions” because of their thin, plate-like structure and apparent function as a newly revealed type of epithelial junction. The lamellar sheets extended for distances as long as a typical cell body but were as thin as 40 nm (micrographs in Figure 4A and three-dimensional rendering in Figure 4F; lamella from cell segmented in purple extends 7.5 μm in the x-y plane and 6.6 μm along the z-axis). Lamellae from different cells formed zones of overlap and sometimes bifurcated to form a tongue-and-groove structure (Figure 4B, inset) or interweaved structures (Figure 4C, F, black mesh highlights area of overlap, dotted outline indicates full area of spread lamellae). We also identified an ear where the lamellar junctions appeared to have been pushed apart (Figure 4D). This may be an ES in the act of deflating via bursting or sliding of the lamellar junction. The full serial EM dataset also confirmed that the endolymphatic duct connects the lumen of the otic vesicle to the tip of the ES, where basal lamellae form a complete barrier (full EM data set available to navigate at <u>http://zebrafish.link/hildebrand16</u>, specific ES links in materials and methods)(Hildebrand et al., in press). Basal lamellae were present along the length of the endolymphatic duct. However, unlike the ES where the apical and lateral membranes separate to expose lamellae, cell bodies in the duct remain tightly packed with apical junctions (Figure 4A,E). Apical junctions in the mutant *lmx1bb^jj410/jj410^* ES remain in contact (Figure 4G) and apparently restrain the pressure without engaging lamellae. This suggests that *lmx1bb*-dependent activity is necessary to expose basal lamellar junctions as the epithelial permeability barrier.

**Figure 4.**
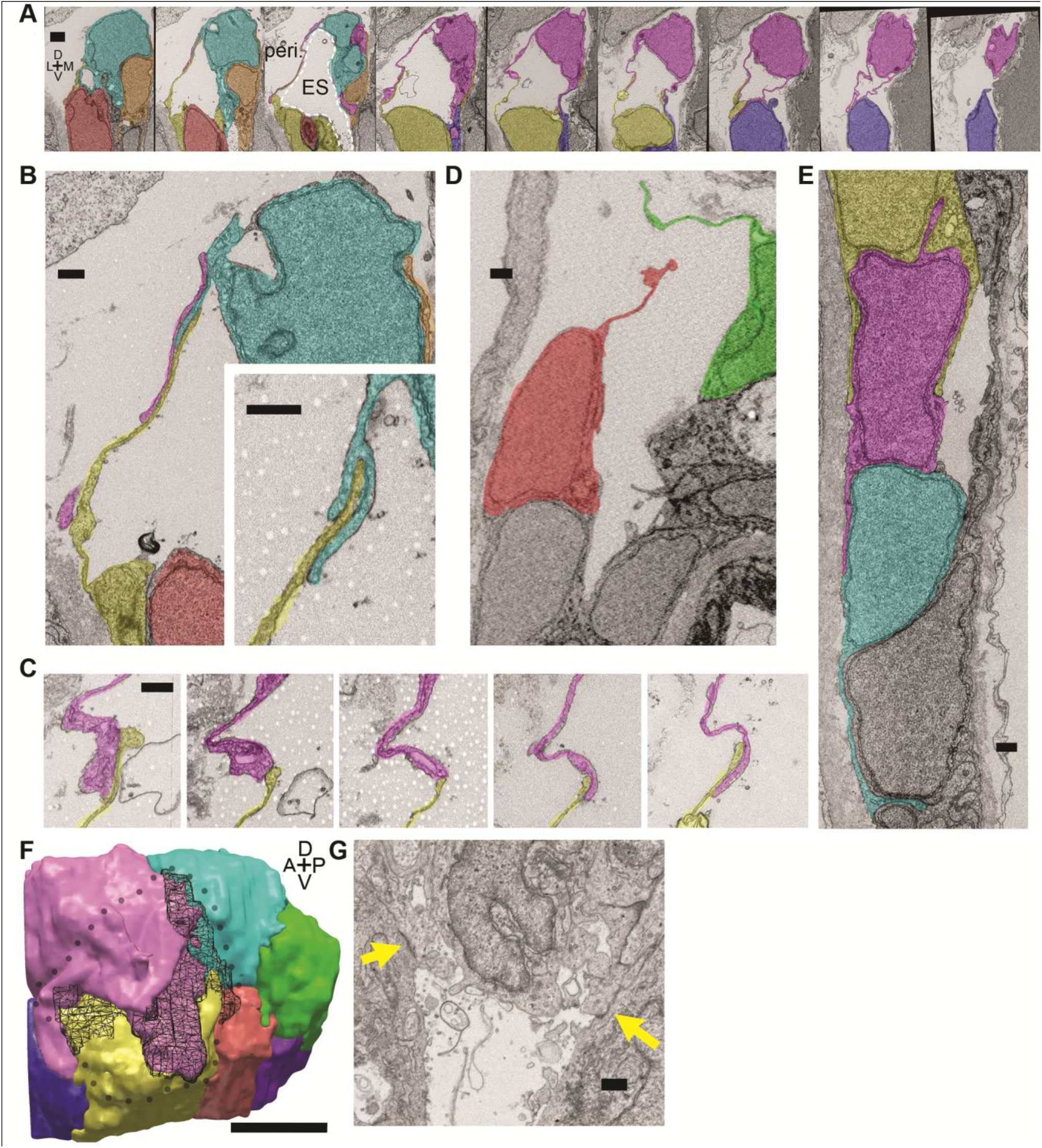
Lamellar junctions at the tip of the ES open to release pressure. (**A**) Select images from serial-section scanning electron microscopy of 5.5 dpf zebrafish. Belonging to a right inner ear, dorsal is up, lateral is left, medial right, ventral down, anterior top, and posterior bottom of the z-stack. Cells forming lamellar junctions were segmented with color overlays to highlight connectivity of lamellae. Presented slices are a subset from the series, each separated by 960 nm (see complete data presented in Video 7). The lumen of the ES lumen is labeled and outlined with a white dashed line in third panel; Perilymph (peri.). (**B**) Lamellae interdigitate and can form tongue-in-groove structures (inset). (**C**) Lamellae can interweave.(**D**) Example of opened lamellar junctions. (**E**) Cells in endolymphatic duct have basal lamellae (presented duct connects with ES of panel A, yellow overlay highlights same cell). (**F**) 3D rendering of ES segmentation from serial micrographs shown in panel (**A**) and Video 7. Black dotted-outline encompasses area of engaged, endolymph filled lamellae. Black mesh highlights area of membrane overlap between lamellae that are spread open. (**G**) *lmx1bb^jj410/jj410^* embryos maintain apical junctions (yellow arrows). Scale bar in (**A**) is 1000 nm, (**F**) is 5 μm, and all other scale bars are 500 nm.

### Cells stretch during ES inflations

Increased pressure could cause an increased ES volume by inducing cell stretching or basal lamellae expansion. To distinguish how the tissue behaves through cycles of ES inflation and deflation we sought better resolution during live imaging. Lattice light-sheet microscopy (LLSM) generates thin light-sheets using Bessel beams to enhance axial resolution and minimize phototoxicity and bleaching (Chen et al., 2014; Gao, Shao, Chen, & Betzig, 2014). Emitted light from the ES passes through brain tissue that scatters and refracts light in a complex, spatially uneven manner. Recent advances in the application of adaptive optics (AO) to microscopy compensate for these uneven aberrations (Wang et al., 2014). A microscope combining live-cell lattice light-sheets and adaptive optics was built (manuscript in preparation) and used here to image ES cycles with significantly improved spatial and temporal resolution.

Because LLSM requires the illumination plane and optical axis to be perpendicular to one another we developed a 3D printed mold for an agarose mount shaped like a volcano that grips the embryo in the desired position (Figure 5A). With proper orientation of the embryo, the LLSM without adaptive optics correction produced good images, which were still a bit blurry with markedly low signal-to-noise (SNR, Figure 5B). The addition of adaptive optics compensated for tissue aberrations and produced images with higher SNR and better contrast (AO-LLSM). The high quality of images enabled us to use our software ACME to reconstruct the membrane signal (Figure 5C), segment cells and the lumen of the ES (Figure 5D, E), and track cells and the lumen accurately through time (Mosaliganti, Noche, Xiong, Swinburne, & Megason, 2012). Similar to the confocal live imaging, we observed cycles of the ES lumen inflating and deflating (Figure 5F, G, Videos 8-9).

**Figure 5.**
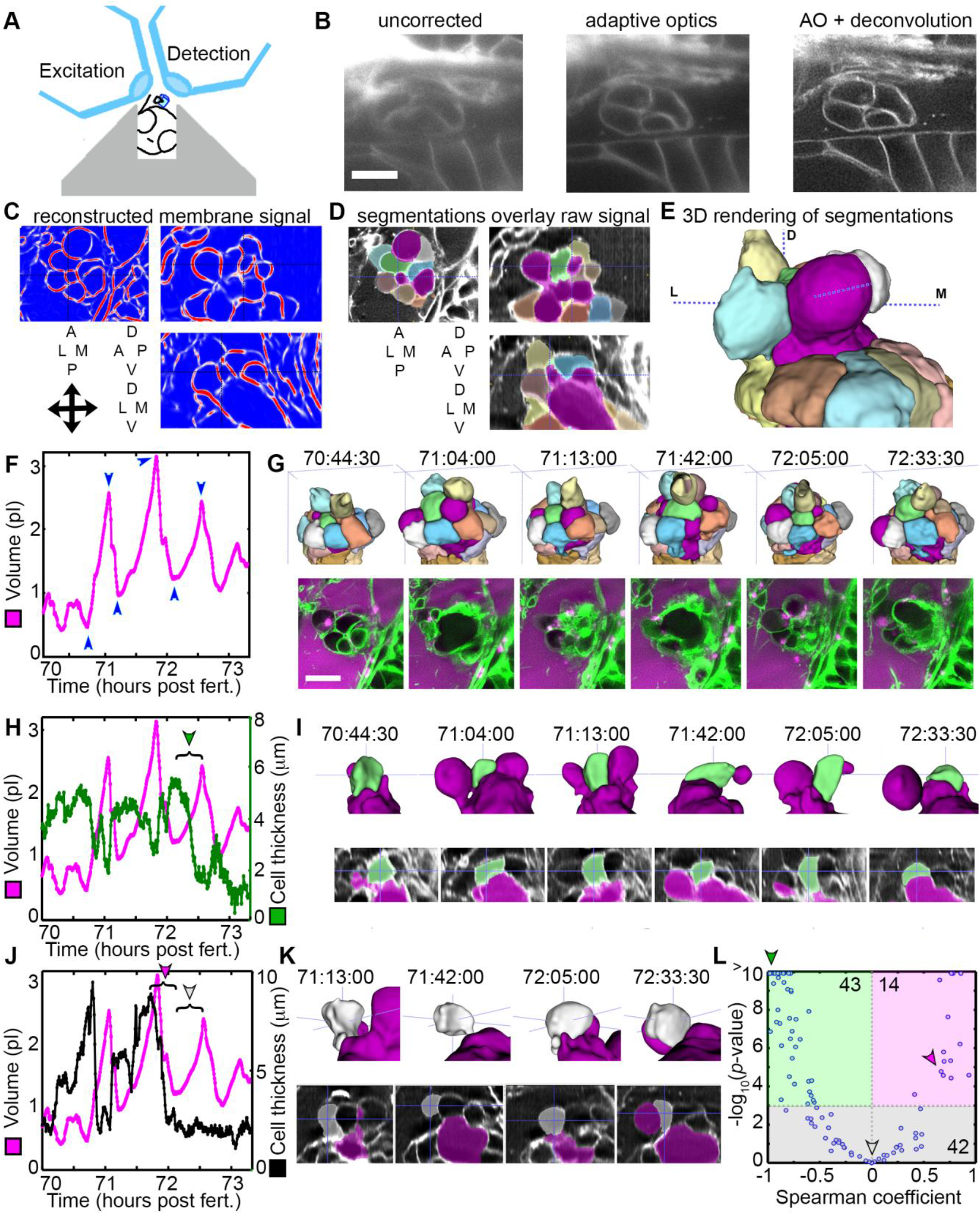
AO-LLSM reveals dynamics of ES cells. (**A**) Illustration of AO-LLSM mounting strategy for imaging ES using volcano mount. (**B**) Representative LLSM images without adaptive optics (AO), with AO, and with AO followed by deconvolution. (**C**) Three orthogonal views of ACME membrane reconstruction. (**D**) Three orthogonal views of raw fluorescence signal overlaid with cell segmentations and ES lumen segmentation (magenta). (**E**) 3D rendering of segmented cells and ES lumen (magenta). (**F**) Volume measurements of segmented ES lumen, imaged every 30 seconds for over 3 hours (Videos 8-10). Blue arrowheads point to time points presented in G and I. (**G**) Top, dorsal perspective displaying 3D renderings of segmented cells. Bottom, maximum intensity projections (MIP) of 4.5 μm slab through tip of ES shows raw data of the ES for the same time points. (**H**) Secondary axis presents green cell’s thickness versus time (green cell in (**G**)). Again, primary axis is volume of ES lumen for comparison. (**I**) Top, 3D renderings of just green cell from (**G**) and magenta lumen highlight stretching of cell, dorsal-medial perspective. Bottom, centered cross-sectional view of raw data overlaid with green cell’s and ES lumen’s segmentations for the same time points. (**J**) Secondary axis is plot of grey cell’s thickness (grey in (**G**)). (**K**) Top, 3D renderings of only grey cell and magenta lumen. Bottom, centered cross-section view of raw data overlaid with grey cell’s and ES lumen’s segmentations for same time points. (**L**) Scatter plot of results from Spearman correlation test of cell thickness trajectories and ES lumen volume trajectories for individual inflation and deflation intervals that are monotonic (example intervals bracketed in H and J). Green region highlights significant correlation (*p*-value < 10^−3^) between cells thinning during inflation or thickening during deflation. Green arrowhead points to test result for bracketed interval in H. Grey region highlights instances where there is no significant correlation between the trajectory of cell thickness and lumen volume. Grey arrowhead points to test result for bracketed interval in J (grey arrowhead). Magenta region highlights significant correlation between cells thinning during deflation or thickening during inflation. Magenta arrowhead points to test result for bracketed interval in J (magenta arrowhead). Y-axis was capped at 10 so that all values greater than or equal to 10 plot as 10. N=99. All scale bars 10 μm.

Cell bodies within the ES stretch in response to increasing pressure. While the increased resolution of AO-LLSM did not allow us to resolve overlapping basal lamellae (thickness ~40 nm), it did allow us to segment the volume of individual cell bodies (lengths and thicknesses ~1-10 μm) and determine their centroids. By measuring the distance between the surface of the ES lumen and each cell body’s centroid, we calculated the apico-basal thickness of each cell. We found that some cells stretched and thinned when the ES lumen was inflated and thickened upon deflation, likely a result of their elastic properties (Figure 5H, I). However, there were instances when the cells at the tip of the ES did not thin during inflation, or while they thinned during some events, they did not thin during others (Figure 5J, K). To quantify these correlations and obtain an overview of the range of behaviors, we determined the Spearman correlation coefficient between trajectories of lumen volume and cell thickness for intervals that spanned individual inflation and deflation events (Figure 5L, bracketed examples in Figure 5H, J). For 43 of 99 tested trajectories, either cell thinning significantly correlated (*p*-value less than 10^−3^) with inflation or cell thickening significantly correlated with deflation (green region, Figure 5L, example of significantly correlated interval, bracket with green arrowhead, Figure 5H, same data point indicated with green arrowhead in Figure 5L). For 42 of 99 tested trajectories, cell behavior did not significantly correlate with inflation or deflation (grey region, Figure 5L, example of uncorrelated inflation interval, bracket with grey arrowhead, Figure 5J, same data point indicated with grey arrowhead in Figure 5L). Unexpectedly, for 14 of 99 tested trajectories there was significant correlation between inflation and cell thickening or deflation and cell thinning (magenta region, Figure 5L, example of correlated interval, bracket with magenta arrowhead, Figure 5J, same data point indicated with magenta arrowhead in Figure 5L). While each cell’s behavior is varied, these data suggest that stretching of a subset of cells contributes to regulation of increased pressure. Additionally, some cells appear to be pushed away from the ES lumen during inflation and pulled back into the ES epithelium during deflation. The complexity of the response is likely the result of features we did not resolve such as the dynamics of each cell’s basal lamellae, each cell’s residual apical and lateral adhesion, and each cell’s basal interface with the extracellular matrix.

### Basal lamellae are dynamic

The low temporal resolution of our original confocal time courses made it unclear whether basal lamellae are static structures like floodgates, only opening to release endolymph, or dynamic. Sparse labeling of cells in the ES further confirmed a ubiquity of lamellae in the ES as well as a variety in their organization (Figure 5A-E, basal side of ES highlighted by dotted line). 3D rendering of the AO-LLSM signal shows that some lamellae are rapidly extending and retracting like they are crawling over neighboring cells or neighboring lamellae (Video 10). The basal lamellae are composed of two juxtaposed membranes and can overlay abutting membranes of adjacent cells or additional lamellae (Figure 4). While we observed stacked membranes by EM, they are still unresolved by AO-LLSM but one might expect patches of increased membrane signal several times the brightness of that of a lone cell membrane. In 3D renderings of the time course we see dynamic patches of increased intensity that can shift rapidly (Figure 6F, Video 10). By merging 3D renderings of 3 consecutive time points, 30 seconds apart, in red, blue, and then green, we observed the relative displacement of these lamellae that can crawl a few microns per minute (Figure 6G, H).

**Figure 6.**
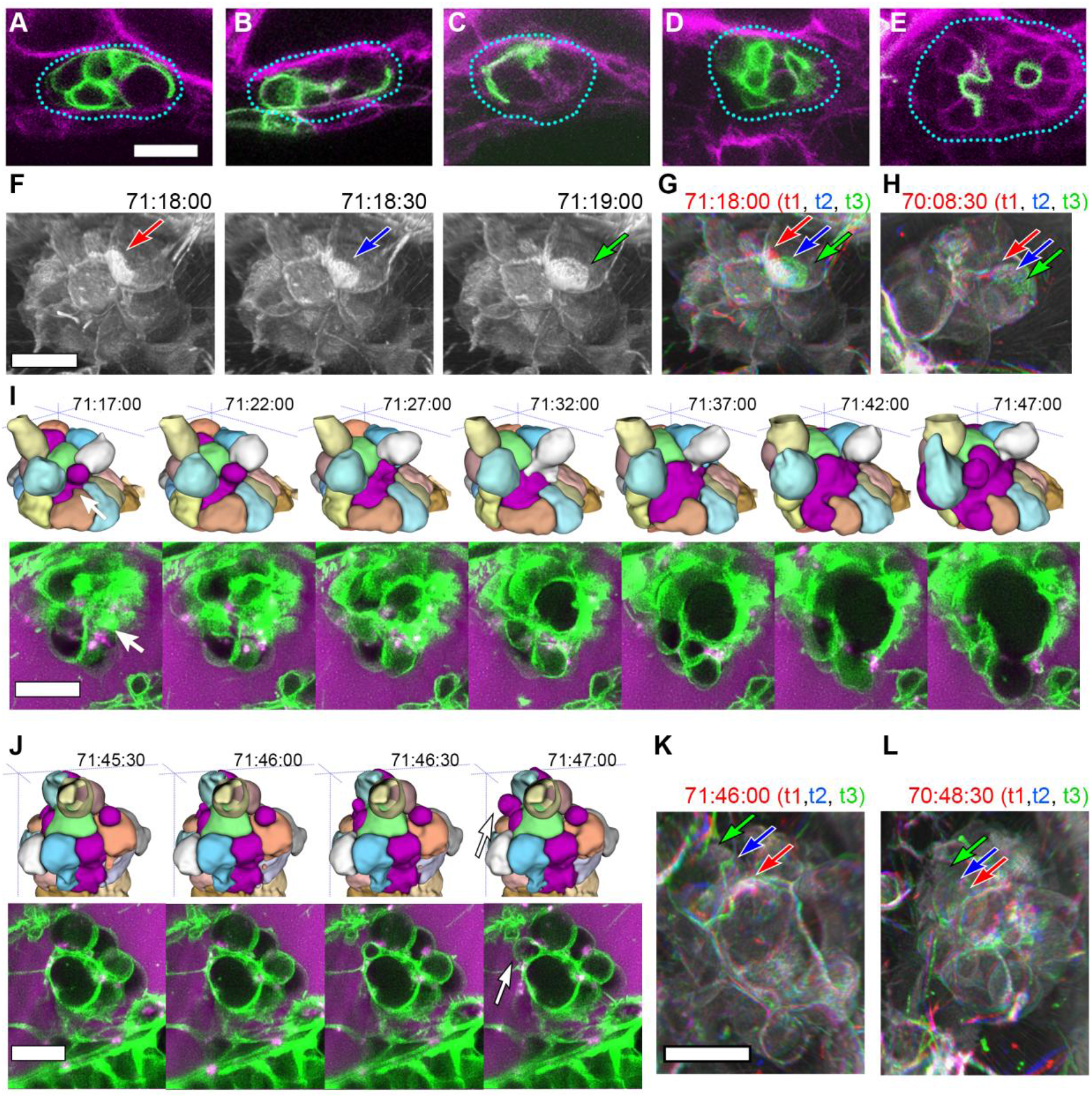
Basal lamellae are dynamic. (**A-E**) Sparsely labeled ES cells: membrane citrine (green) in a membrane cherry background (magenta). Blue dotted line traces basal sides of ES tissue. (**F**) 3D rendering of AO-LLSM data, 3 sequential time points 30 seconds apart (membrane citrine depicted in grey, Video 10). Bright patches on surface move (red, blue, green arrows). (**G**) Consecutive 3D renderings overlaid as red (71:18:00), blue (71:18:30), and then green (71:19:00). Immobile regions remain grey, while regions of displacement are red, blue, and green. Arrows point to moving lamellae. (**H**) Additional example of moving lamellae, again visualized by overlaying consecutive images in red (70:08:30), blue (70:09:00), and then green (70:09:30). Arrows point to moving lamellae. (**I**) Top, time points of 3D rendered segmentations spanning 30 minutes. Below, 4.5 μm MIP slabs of the raw data for the same time points. For both views, arrow points to same region where lumen segmentation is slowly exposed as thin lamellar region expands. (**J**) Top, time points of 3D rendered segmentations spanning 2 minutes. Below, 4.5 μm MIP slabs of the raw data for the same time points. For both views, arrow points to same region where lumen segmentation rapidly inflates as thin lamellar region expands. (**K**) Consecutive 3D renderings, same as (**J**), overlaid as red (71:46:00), blue (71:46:30), and then green (71:47:00). Arrows point to rapidly inflating lamellae. (L) Consecutive 3D renderings, overlaid as red (70:48:30), blue (70:49:00), and then green (71:49:30). Arrows point to rapidly inflating lamellae, same region as (K). All scale bars 10 μm.

While the resolution of AO-LLSM is improved, it does not resolve the double membrane structure of lamellae or enable identification of the cell from which a lamella extends. Thus, in our surface-rendered segmentations, we give the cell membranes of each cell body a unique pseudo-color while the lamellae that are just exposed to the lumen and not adjacent to another cell body are collectively colored magenta (Figure 5E, G, corresponds to black dotted outline in Figure 4F). By visualizing the segmented objects in 3D we can identify when lamellar junctions are exposed based on when and where the magenta appears (Figure 5G, 6I, Video 9). Inflations of the ES lumen coincided with slow increases in the surface area of exposed lamellae (30 minute expansion, Figure 6I). The expansion of thin membrane signal was also observable in the raw data (bottom portion of panel, Figure 6I). Some secondary loci included smaller lamellae that engaged and inflated much more quickly (1-2 minute expansion, Figure 6J). We can visualize their sudden expansion by again merging consecutive time points, 30 seconds apart, with red, blue, and then green (Figure 6K). This presentation reveals that the same locus can rapidly expand multiple times (one minute rapid expansions an hour apart, Figure 6K,L). In summary, multiple sets of basal lamellae can simultaneously engage at different loci, either through slow spreading or rapid expansion.

Given the diversity and speed of lamellar dynamics we asked whether Rac1, known to promote actin polymerization in lamellipodial protrusions, was important for ES valve function (Waterman-Storer, Worthylake, Liu, Burridge, & Salmon, 1999). Heat-shock induction, throughout the entire embryo, of a dominant negative Rac1 at 56 hpf resulted in otic vesicles that became leaky and subsequently collapsed between 60-70 hpf (Video 11, 19/34 leaky and collapsed ears versus 1/12 in a heat-shock gfp control)(Kardash et al., 2010). Notwithstanding slower development likely caused by the many roles of Rac1, heat shock induction of the dominant negative Rac1 at 32 hpf, prior to ES formation, did not result in leaking or ear collapse during the subsequent 15 hours (while 0/20 leaky or collapsed ears versus and 0/14 in a heat-shock gfp control). These results suggest that the basal lamellae in the ES may utilize cytoskeletal regulators also used for lamellipodial extension.

### Basal lamellae open for deflation

While we had evidence that lamellae can separate in our EM images and evidence that the epithelial barrier breaks during deflation, as seen in the leak in fluorescence from labeled perilymph, it remained uncertain whether separation of lamellae precipitated the deflation events. To observe evidence of the lamellar valve opening, we looked for instances when the membrane signal of lamellar junctions was disrupted prior to deflation events. We were able to witness instances when membrane signal from inflated lamellae was disrupted prior to deflation (arrows, Figure 7A-D). These events can be accompanied by a thin lamellar protrusion sticking out into the perilymph (arrow, Figure 7D). Examination of the raw and rendered segmentations prior to and following breaks in membrane signal showed that it preceded a deflation event (reduction in local volume of ES lumen, magenta, Figure 7A,C).

**Figure 7.**
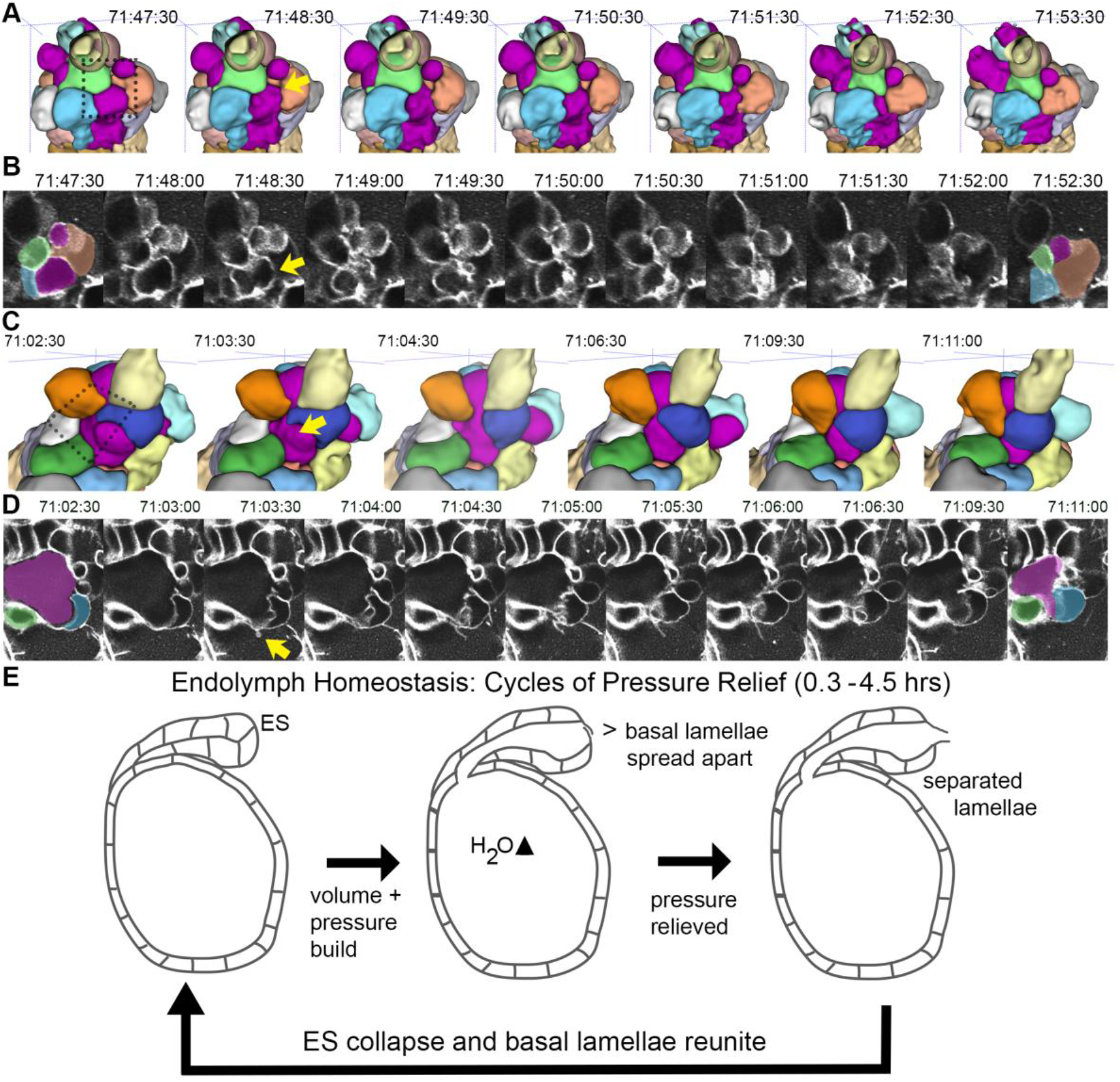
Basal lamellae open prior to deflation. (**A,C**) 3D renderings of segmented cells and ES lumen. (**A**) Dorsal view. (**B**) Raw data of membrane citrine AO-LLSM data spanning same time range as in (**A**). First and last time point are overlaid with segmentations for cells and lumen (magenta) neighboring the region of valve opening. In (**A-D**) arrows point to site of lamellae separating, dotted outlines indicate field of views in B and D. (**C**) Anterior view of another instance of lamellae separating. (**D**) Raw data spanning time range of (**C**). (**E**) Illustrated pressure relief mechanism.

## DISCUSSION

We report identification of the ES as a pressure relief valve based on six lines of evidence. First, the lumen of the ES slowly inflates and then rapidly deflates every ~1–4 hours. Second, deflation of the ES coincides with a breach in its epithelial diffusion barrier. Third, laser ablation of the otic epithelium is sufficient to induce rapid deflation of the inflating ES. Fourth, the ultrastructure of the closed and opened relief valves in the ES shows that lamellar junctions can pull apart. Fifth, high temporal and spatial resolution time courses reveal lamellae that behave like lamellipodia: constantly crawling over one another before they separate to relieve pressure and volume. Sixth, perturbation of the valve’s development with a genetic mutation causes distension of the ES as found in common ear disorders such as Meniere’s disease, Pendred syndrome, and Enlarged Vestibular Aqueduct syndrome. We conclude that increased pressure is managed through a combination of strain that stretches viscoelastic cells in the ES and adhesion distributed across the surface area of dynamic basal lamellae that transiently provides a barrier before they open to release pressure (Figure 7E). Herein, we discuss the data and its limits as well as our main conclusion with respect to the physiology and pathology of the inner ear.

A physical relief valve in the ES, composed of overlapping basal lamellae, had not been previously identified because of the lack of sufficient temporal and spatial resolution as well as the lack of a system with an optically accessible ES. Thin overlapping basal junctions are seen at the marginal folds of lymph and blood capillaries (Baluk et al., 2007). Capillaries flank the developing and adult ES and the similarity between the cross-sectional ultrastructure of the ES’s lamellar junctions and capillaries, as well as vacuoles, could have led to inaccurate annotation of lamellar junctions (Kronenberg & Leventon, 1986). EM micrographs of the rat, guinea pig, tree frog, and human ES showed instances of thin cytoplasmic extensions enclosing large lumens, which were annotated as capillaries or large vacuoles (Bagger-Sjoback, Friberg, & Rask-Anderson, 1986; Bagger-Sjoback & Rask-Andersen, 1986; Dahlmann & von During, 1995; Kawamata, Takaya, & Yoshida, 1987; Moller, Caye-Thomasen, & Qvortrup, 2013; Qvortrup, Rostgaard, Holstein-Rathlou, & Bretlau, 1999). More recent EM studies of the human ES, which used more refined techniques to prepare samples from adult cadavers, recognized similar structures as belonging to the distal portion of the ES and described them as dilated tubili with an interconnected lumen resembling a network of cisterns (Moller et al., 2013). While appearing more elaborate in humans, the structural unit of the relief valve seems to be conserved. However, because of insufficient spatial and temporal resolution, it remains to be determined whether these cells have overlapping lamellae and whether they function as a physical relief valve.

Primarily, pressure relief prevents endolymphatic hydrops—the buildup of pressure and potential tearing of the ear’s epithelium—that is associated with many inner ear pathologies. Secondarily, low-frequency deflations of the ES may drive intermittent longitudinal flow of endolymph from the cochlea and semicircular canals into the endolymphatic duct and sac. The presence of longitudinal flow had been proposed to explain the accumulation of debris and tracers within the ES and, while never directly observed in an unperturbed inner ear, longitudinal flow can be induced and observed through manipulations of the inner ear (Salt, 2001; Salt & DeMott, 1998; Salt & Plontke, 2010; Salt, Thalmann, Marcus, & Bohne, 1986). The localized inflation and patterned formation of valve cells leads to a localized break at the ES lamellar junctions. The local break could create a transient pressure gradient that leads to flow from other parts of the inner ear towards the endolymphatic duct and sac. The intermittent and slow nature of this flow could explain conflicting interpretations of classic studies using tracer injections where tracer accumulated in the ES after long periods of time but significant flow was absent when observed on short time scales (Guild, 1927; Manni & Kuijpers, 1987; Salt, 2001; Salt et al., 1986).

Lmx1bb is a LIM homeobox transcription factor. Mutations in *LMX1B* cause Nail Patella syndrome in humans and a third of Nail Patella syndrome patients develop glaucoma, a phenotype also observed in mouse models (Cross et al., 2014; Liu & Johnson, 2010; Mimiwati et al., 2006). The mechanism by which glaucoma arises in the disease and in mouse models is not clear; however, the trabecular meshwork and Schlemm’s canal appear abnormal in these cases. Giant vacuoles and pores in Schlemm’s canal may be used to relieve intraocular pressure, and their ultrastructure resembles the lamellar junctions of the ES (Gong, Ruberti, Overby, Johnson, & Freddo, 2002). Like the ear, pulsatile pressure relief occurs in the eye, although at the higher frequency of the heart rate, and Schlemm’s canal is a major site of the pressure relief (Ascher, 1961; Johnstone, Martin, & Jamil, 2011). Additionally, up to half of Nail Patella syndrome patients suffer from kidney disease(Bennett et al., 1973). During podocyte maturation apical junctions are lost, thereby enabling development of the basal slit diaphragm that filters blood(Pavenstadt, Kriz, & Kretzler, 2003). In mice lacking LMX1B activity, podocytes fail to lose their apical junctions(Miner et al., 2002). Removal of apical junctions may thus be a specialized mechanism used in organs where elaborations of junction architectures allow controlled release of fluids from one compartment into another. We speculate that LMX1B may have a common set of targets in the ES, eye, and kidney involved in regulated apical junction remodeling.

We find that basal lamellae in the endolymphatic sac behave like a pressure relief valve to regulate fluctuating volume and pressure that arises from excess endolymph in the inner ear (Figure 7E). The high temporal and spatial resolution of AO-LLSM, combined with image processing tools for segmenting, tracking, and quantifying cell geometries, was necessary to reveal the dynamic cell behaviors underlying the pulsing of the ES. While cells of the ES are immobile epithelial cells, the cell extensions resemble lamellipodia of crawling cells in thickness (40 nm), speed (as fast as ~1 μm per minute), and regulatory mechanism (Rac1)(Abercrombie, 1980). Because of limitations of imaging, it remains unclear how each cell’s basal lamellae, residual apical and lateral adhesion, and basal interface with the extracellular matrix contribute to the physical behavior of the ES. Of related interest is how the crawling lamellae adhere to their substrate in the ES to generate a barrier without gaps during the inflation phase. The force of this adhesion, in combination with the viscoelastic properties of the ES, likely determines the relief valve’s set point for volume and pressure homeostasis. For the inner ears of adult mammals, this set point is likely around 100-400 Pascals, although it is unknown how much the inner ear’s pressure and volume fluctuate (Park et al., 2012). Determination of the molecular basis of lamellar junction organization will be necessary to make progress on how the physical relief valve works and how it might fail to cause disease.

## Materials and methods

### Zebrafish strains and maintenance

Zebrafish were maintained at 28.5°C using standard protocols (Westerfield, 1993). The Harvard Medical Area Standing Committee on Animals approved zebrafish work under protocol number 04487. Adult zebrafish, 3 months to 2 years of age, were mated to produce embryos and larvae. Adults were housed in a main fish facility containing a 19-rack Aquaneering system (San Diego, CA) and a 5-rack quarantine facility for strains entering the system from outside labs or stock centers. The systems’ lights turn on at 9am and go off at 11pm. Fish matings were set up the night before with males separated from females with a divider in false-bottom crossing cages in a pair wise, or 2×2 fashion to maximize embryo yield. The divider was pulled the following morning, shortly after the lights turned on. Egg production was monitored to establish the time of fertilization. Manipulations and observations of baby zebrafish were performed between fertilization and 5.5 days post fertilization.

These studies were performed using the AB wild-type strain, the *lmx1bb^jj410/jj410^* mutant and the following transgenic lines: *Tg(actb2:mem-citrine-citrine)^hm30^, Tg(actb2:mem-citrine)/(actb2:Hsa.H2b-tdTomato)^hm32^, Tg(actb2:mem-citrine)/(actb2:Hsa.H2b-tdTomato)^hm33^* (these three alleles were combined for maximal membrane signal, *hm32* and *hm33* are two separate alleles of the same construct that is composed of two divergent beta-actin promoters, one driving membrane citrine and the other driving histone tdTomato, *actb2:Hsa.H2b-tdTomato* of these divergent constructs tends to be silenced in transgenic fish and is not useful), *Tg(−5.0lmx1bb:d2eEGFP)^mw10^, Tg(actb2:mem-mcherry2)^hm29^, Tg(hsp70:rac1_T17N-p2a-mem-cherry2)^hm35^*, and *Tg(elavl3:GCaMP5G)^a4598^* (Ahrens, Orger, Robson, Li, & Keller, 2013; McMahon, Gestri, Wilson, & Link, 2009; Obholzer et al., 2012; Schibler & Malicki, 2007; Xiong et al., 2013).

### Immobilization and time-lapse imaging

Embryos were immobilized with either 50 pg of α-bungarotoxin mRNA injected into the 1-cell embryo or 2.8 ng of α-bungarotoxin protein injected into the heart at 59 hpf (Swinburne et al., 2015). α-bungarotoxin mRNA was synthesized from a linearized plasmid using the mMessage mMachine T7 ULTRA kit (Thermo Fisher Scientific, Waltham, MA). Subsequently, mRNA was purified using RNAeasy Mini Kit (Qiagen, Hilden, Germany). α-bungarotoxin protein was obtained from Tocris Bioscience (Bristol, United Kingdom). 2.3 nL injections were performed using Nanoject II (Drummond Scientific, Broomall, PA). For tracking perilymph, ~10 ng of 3 kDa dextran-Texas red neutral (Thermo Fischer Scientific) was injected into the zebrafish heart at 59 hpf. Immobilization with α-bungarotoxin permits the ear to grow and develop normally (Swinburne et al., 2015).

Paralyzed embryos were cultured in Danieau buffer and mounted in an immersed 1.5% w/v agarose canyon mount or the volcano mount for AO-LLSM, that were generated from custom-made molds. The canyons were 0.4 mm wide and 1.5 mm deep. The volcano mold for AO-LLSM was printed by shapeways.com (New York City, NY). The inner width of the volcano was also 0.4 mm and was printed using their “Frosted Extreme Detail” material. The embryo was placed dorso-laterally in the canyon or volcano so that dorsal portion of the left otic vesicle was either flush with a #1 coverslip placed over the canyon (embryos were tilted approximately 30 degrees laterally from their dorsal axes) or protruding from the mouth of the volcano mold. No coverslip was placed above embryos for AO-LLSM. Homemade hair loops were used to position embryos.

Mounted embryos were imaged on an upright LSM 710 (Carl ZEISS, Göttingen, Germany) with a C-apochromat 40x / NA 1.2 objective (ZEISS). The objective’s corrective ring was adjusted to account for the use of #1 coverslips (0.13-0.16 mm thick). Imaging took place within a homemade foam-core incubator maintained at 28.5°C. Lasers of wavelength 405, 488, 514, and 594 nm were used to image GFP, citrine, Texas red, and mCherry2. A typical imaging session used the following set up: citrine excited with 514 nm laser 5% (Ch1 filters 519-584 nm, gain 885), Texas red excited with 594 nm laser 15% (Ch2 filters 599-690 nm, gain 864), 1.27 μs pixel dwell, 1 line average, 137 μm pinhole, 1.2 zoom, 458/514/594 beam splitter, and 0.173 × 0.173 × 1.135 μm voxel scaling. Around 20-30 long-term time courses were attempted to establish the method. The data presented is representative of 8 wild-type time courses and 4 mutant time courses. Additionally, hundreds of mutants were observed with distended ES’s during the cloning of the mutant and phenotype characterization at 72 and 96 hpf.

Laser ablations of epithelial cells in the otic vesicle were performed using a similar imaging set-up as for time-lapse confocal imaging. For the cell ablations a Mai-Tai HP 2-photon laser (Spectra-Physics, Santa Clara, CA) was used. After a target was chosen the 2-photon laser was tuned to 800 nm at 50% power, the pinhole was opened completely, the 690+ beam splitter was selected, and a spot scan was performed for 10,000 cycles. 2 or 3 spots were targeted in adjacent cells to ensure the wounds disrupted the epithelial barrier. After ablations, time-lapse microscopy was performed with 1-photon microscopy, as described above. The data presented is representative of 4 such experiments as well as 7 ablations that were evaluated without time courses, 1 hour after ablation.

For imaging with the adaptive optics lattice light-sheet microscope, zebrafish immobilized with α-bungarotoxin were mounted in a 1.5% low melt agarose volcano mold on a 5 mm coverslip and imaged starting at 68-72 hpf. We submerged the excitation and detection objectives along with the 5 mm coverslip in ~8 ml of 1X Danieau buffer at room temperature (22°C ±2°C). The zebrafish tissues expressing membrane citrine marker and 3 kDa dextran Texas Red fluid phase marker were excited using 488 nm and 560 nm lasers (488 nm operating at ~3 mW and 560 nm operating at 5 mW corresponding to ~16 μW and ~31 μW at the back aperture of the excitation objective) both sequentially exposed for 15 msec. The tissues were excited with 488 nm and 560 nm sequentially by dithering a multi-Bessel light-sheet arranged in a square lattice configuration (corresponding to an excitation inner/outer numerical aperture of 0.517/0.55, respectively). The optical sections were collected by scanning the detection objective with 300 nm steps. Each imaging volume consisted of 131 optical planes capturing 1024 × 1024 pixels, thereby capturing a volume of ~ 97 μm × 97 μm × 39 μm every 15.5 or 29.9 seconds using a dual Hamamatsu ORCA-Flash 4.0 sCMOS camera setup (Hamamatsu Photonics, Hamamatsu City, Japan). Prior to the acquisition of the time series data consisting of 300-530 time points, the imaged volume was corrected for optical aberrations (manuscript in preparation). The imaged volumes were deconvolved with experimentally measured point-spread functions measured with 0.17μm tetraspec beads (Thermo Fisher) excited with 488 nm and 560 nm lasers in MATLAB using Lucy-Richardson algorithm on HHMI Janelia Research Campus’ computing cluster and locally with a 3.3 GHz 32-Core Intel Xeon E5 with 512 GB memory. The LLSM was operated using a custom LabVIEW software (National Instruments, Woburn,

MA) on a 3.47 GHz Intel Xeon X5690 workstation with 96 GB memory running Microsoft Windows 7 operating system. The images presented are from one AO-LLSM time course that is representative of three other similar acquisitions also obtained using AO-LLSM.

### Serial-section electron microscopy

“Wild-type” *Tg(elavl3:GCaMP5G)^a4598^* and “mutant” *lmx1bb^jj410/jj410^* larvae at 5.5 days post-fertilization (or 4.5 days for the mutant that begins to suffer from pleiotropic toxicity) were briefly anesthetized with 0.02% tricaine methanesulfonate (Sigma-Aldrich, St. Louis, MO), and subsequently preserved by dissection and immersion into an aldehyde fixative solution (2% paraformaldehyde and 2.5% glutaraldehyde in 0.08M Sorenson’s phosphate buffer, Electron Microscopy Sciences, Hatfield, PA). Each specimen was then post-fixed and stained *en bloc* with reduced osmium solution (1% osmium tetroxide (Electron Microscopy Sciences) and 1.5% potassium ferricyanide(Sigma Aldrich)) followed by uranyl acetate (1% uranyl acetate (Electron Microscopy Sciences) in 0.05 M maleate buffer). Finally, samples were dehydrated with serial dilutions of acetonitrile in distilled water, infiltrated with serial dilutions of epoxy resin in acetonitrile (Electron Microscopy Sciences), and embedded in low-viscosity resin (Koehler & Bullivant, 1973). The epoxy resin for infiltrations and embedding was composed of 63% nonenyl succinic anhydride (Electron Microscopy Sciences), 35.5% 1,2,7,8-diepoxyoctane (97%, Sigma), and 1.5% 2,4,6-tri(dimethylaminomethyl) phenol (DMP-30, Electron Microscopy Sciences), v/v.

Serial-sections through the endolymphatic sac were continuously cut with a nominal thickness of 60 nm using a 45° diamond knife (Diatome, Biel, Switzerland) affixed to an ultramicrotome (Leica EM UC6, Leica Microsystems, Wetzlar, Germany) and collected with an automated tape-collecting ultramicrotome(Hayworth et al., 2014). Field emission scanning EM of back-scattered electrons was conducted either on a Zeiss Merlin (ZEISS) or a FEI Magellan XHR 400L (Thermo Fisher Scientific) with an accelerating voltage of 5.0 kV and beam current of 1.6-7 nA(Hildebrand et al., in press). Image registration was performed with Fiji TrakEM2 alignment plug-ins(Saalfeld, Cardona, Hartenstein, & Tomancak, 2010; Saalfeld, Fetter, Cardona, & Tomancak, 2012; Schindelin et al., 2012). Manual image segmentation was performed with ITK-SNAP(Yushkevich et al., 2006). The data presented is from a single wild-type and single mutant sample because of the time and effort required technology development as well as in sample preparation, acquisition, and processing.

### Quantification and statistical analyses

Time-lapse confocal data sets were converted from Zeiss’s LSM format to a series of image files (*.png) with a header file containing information on the imaging set up (*.meg). This was performed with a script called *lsmtomegacapture* available in our open source “in toto image analysis tool-kit” (ITIAT, <u>https://wiki.med.harvard.edu/SysBio/Megason/GoFigureImageAnalysis</u>). The image data sets were then loaded into GoFigure2, open-source software developed in the Megason lab for image analyses with a database (<u>http://www.gofigure2.org</u>; Gelas, Mosaliganti, and Megason et al., in preparation). In GoFigure2 the ES was manually contoured and these contours were exported as XML-files in the GFX format. Exported contours were modeled in 3D and volumes were quantified using a script called *GoFigure2ContoursToMeshes*, which uses the power crust reconstruction algorithm(Amenta, 2001). 3D meshes were generated in the VTK format and viewed using ParaView (KitWare)(Henderson, 2007). Leak in fluorescence was measured by placing a sphere with radius 1 μm into the lumen of the ES and GoFigure2 recorded the integrated fluorescence intensity.

Automated image analysis of the AO-LLSM data was performed using modified versions of the ACME software, python shell scripts, and ITIAT scripts (Mosaliganti et al., 2012). Starting with 3D tif files, the ACME pipeline includes the following steps, parameters used in (**bold**):

1. *convertFormat inputFile.tif outputFile.mha spacingX(**0.097**) spacingY(**0.097**) spacingZ(**0.3**)*
2. *cellPreprocess inputFile.mha outputFile.mha cellRadius (**0.1**)*
3. *resample inputFile.mha outputFile.mha spacingFactorX(**1.5**) spacingFactorY(**1.5**) spacingFactorY(**0.5**)*
4. *multiscalePlanarityAndVoting3D inputFile.mha outputFile.mha iSigmaFiltering(**0.25**) iSigmaVoting(**0.5**) alpha(**0.3**) beta(**0.25**) gamma(**1000**)*
5. *membraneSegmentation inputResample.mha inputTensorVoting.mha outputSegmentedLabelImage.mha threshold(**1.0**)* Correcting of segmentations and tracking were done using iterative implementations of the ITIAT scripts *membraneSegmentationWithMarkersImageFilter* and *MorpholigcalErosionLabelImageFilter* with manual error corrections done in ITK-SNAP(Yushkevich et al., 2006).

3D rendering of segmented labels and overlays of segmented labels and raw membrane signal were performed using the software ITK-SNAP (<u>www.itksnap.org</u>, (Yushkevich et al., 2006)). 3D visualization of raw membrane data was performed using the software FluoRender (<u>www.sci.utah.edu/software/fluorender.html</u>, (Wan, Otsuna, Chien, & Hansen, 2012)). Videos were compiled and encoded using a combination of ImageJ and HandBrake ((Schindelin et al., 2012), <u>www.handbrake.fr</u>).

For tracking cell thickness, we used the ITIAT scripts *CellSegmentationStatistics* to map the coordinates of each cell’s centroid, *sizeThresh* to extract label image files of just the ES lumen, and *DistanceFromMask* to calculate the physical distance from cell centroids to the surface of the ES lumen label. Cell and lumen volumes were also determined using *CellSegmentationStatistics*.

The outputs from *DistanceFromMask* and *CellSegmentationStatistics* were loaded into MATLAB for both graphing and further analysis. For determining the Spearman correlation coefficient between lumen volumes we first parsed the data matrix into time windows when lumen volume exhibited approximately monotonic increasing or decreasing values. For these time windows we used MATLAB’s statistical toolbox to determine the Spearman correlation coefficient and its associate *p*-value. The syntax for this analysis was: [RHO,PVAL] = corr(LumenVolume,CellThickness,‘type’,‘Spearman’). This test was performed on 99 data windows. MATLAB was also used to perform a Mann-Whitney-Wilcoxon test on the significance cell thickness differences between wild-type and mutant images (Figure 3).

## Data and software availability

The image processing scripts described above are available through GITHUB at the following repositories:

<u>https://github.com/krm15/ACME/tree/MultithreadLookup</u>

<u>https://github.com/krm15/AO-LLSM</u>

<u>https://github.com/krm15/GF2Exchange</u>

The full data-set of serial-section EM images are publicly available at resolutions of 4.0 × 4.0 × 60 nm^3^ per voxel for the closed valve and 18.8 × 18.8 × 60 nm^3^ per voxel for the open valve. To examine the ES anatomy viewers should navigate to <u>http://zebrafish.link/hildebrand16/data/es_closed</u>(closed valve) or <u>http://zebrafish.link/hildebrand16/data/es_open</u>(open valve). This set of multiresolution serial-section EM image volumes was co-registered(Wetzel et al., 2016) to allow one to navigate the lumen of the endolymphatic duct from the lumen of the otic vesicle to the tip of the ES.

## Acknowledgements

We thank Dante D’India for fish care. Brian Link, Erez Raz, Elizabeth Benecchi, and Margaret Coughlin for reagents. Lisa Goodrich, Becky Ward, Amelia Green, Akankshi Munjal, Sandra Swinburne, and the Megason lab for comments. Kerry Sobieski, Betzig lab members, and members of the Janelia Research Campus aquatics facility for helping with travel, shipping, and assisting with experiments at Janelia Research Campus. I.A.S. was supported by NIH fellowship 5F32HL097599, a Hearing Health Foundation Emerging Research Grant, and a Novartis Fellowship in Systems Biology. K.R.M. was supported by NIH grant K25 HD071969. D.G.C.H. was supported by NIH grants T32 MH20017 and T32 HL007901 and NSF IIA EAPSI award 1317014. T.K. acknowledges support from the Janelia Visitor Program, HHMI, Biogen and NIH grant R01 GM075252. S. U. is a Fellow at the Image and Data Analysis Core at Harvard Medical School. This work was supported by R01 DC010791 and R01 DC015478 from the National Institute of Deafness and Other Communication Disorders (S.G.M.) and by NIH grants DP1 NS082121, RC2 NS069407 from the NINDS (F.E.).

---

**Figure 1—figure supplement 1.**
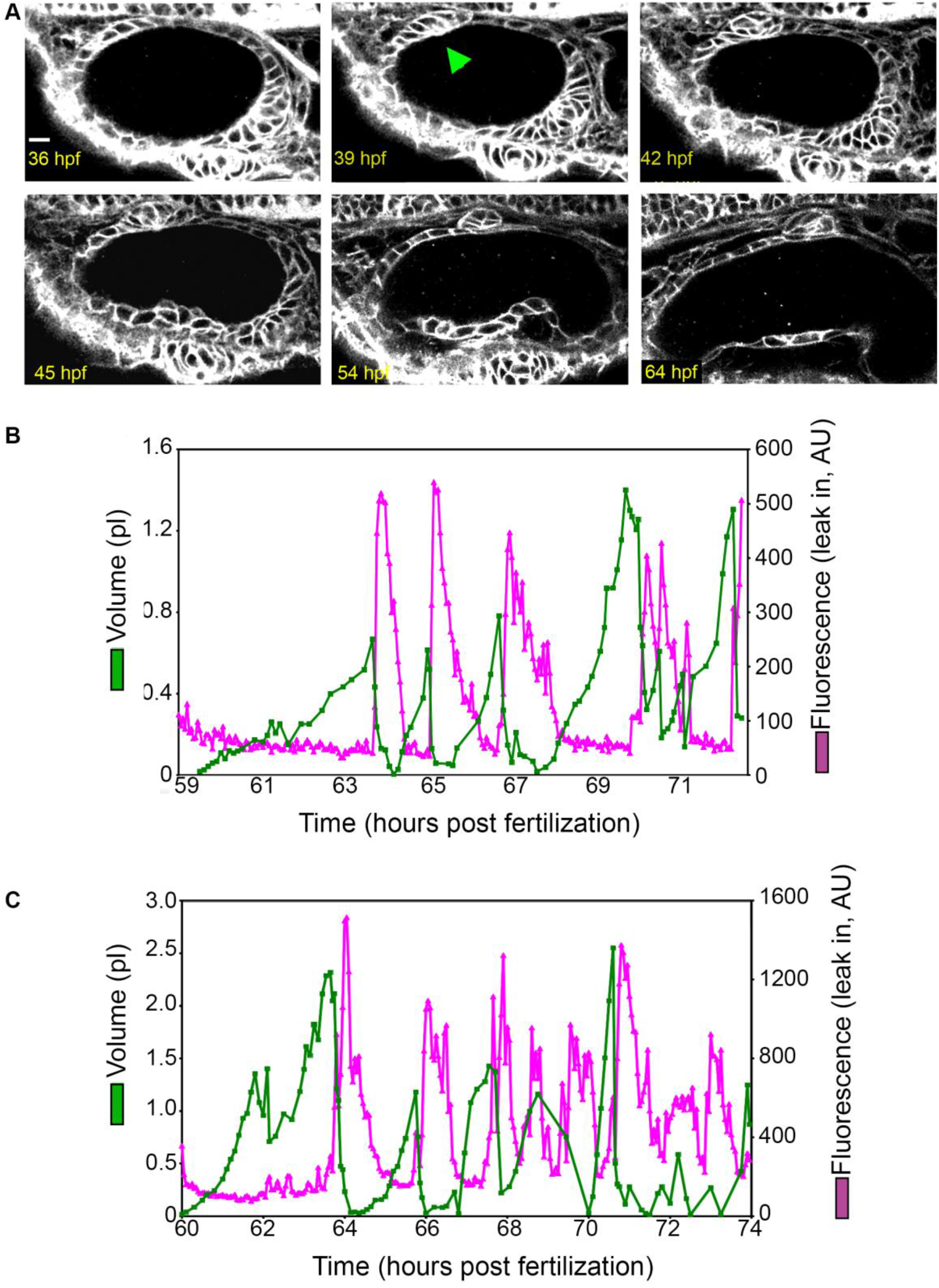
Early ES development and additional examples of wild-type ES inflation and deflation. (A) ES morphogenesis begins at 36 hours post fertilization (hpf) as an evagination in the dorsal epithelial wall of the otic vesicle (green arrowhead points to nascent ES, see also Video 1). Scale bar 10 μm. (B-C) Quantification of segmented ES volumes (primary axis, green) and leak in fluorescence (secondary axis, magenta) over multiple cycles. (B) Analysis of data presented in left panel of Video 3. (C) Analysis of data presented in right panel Video 3.

**Figure 3—figure supplement 1.**
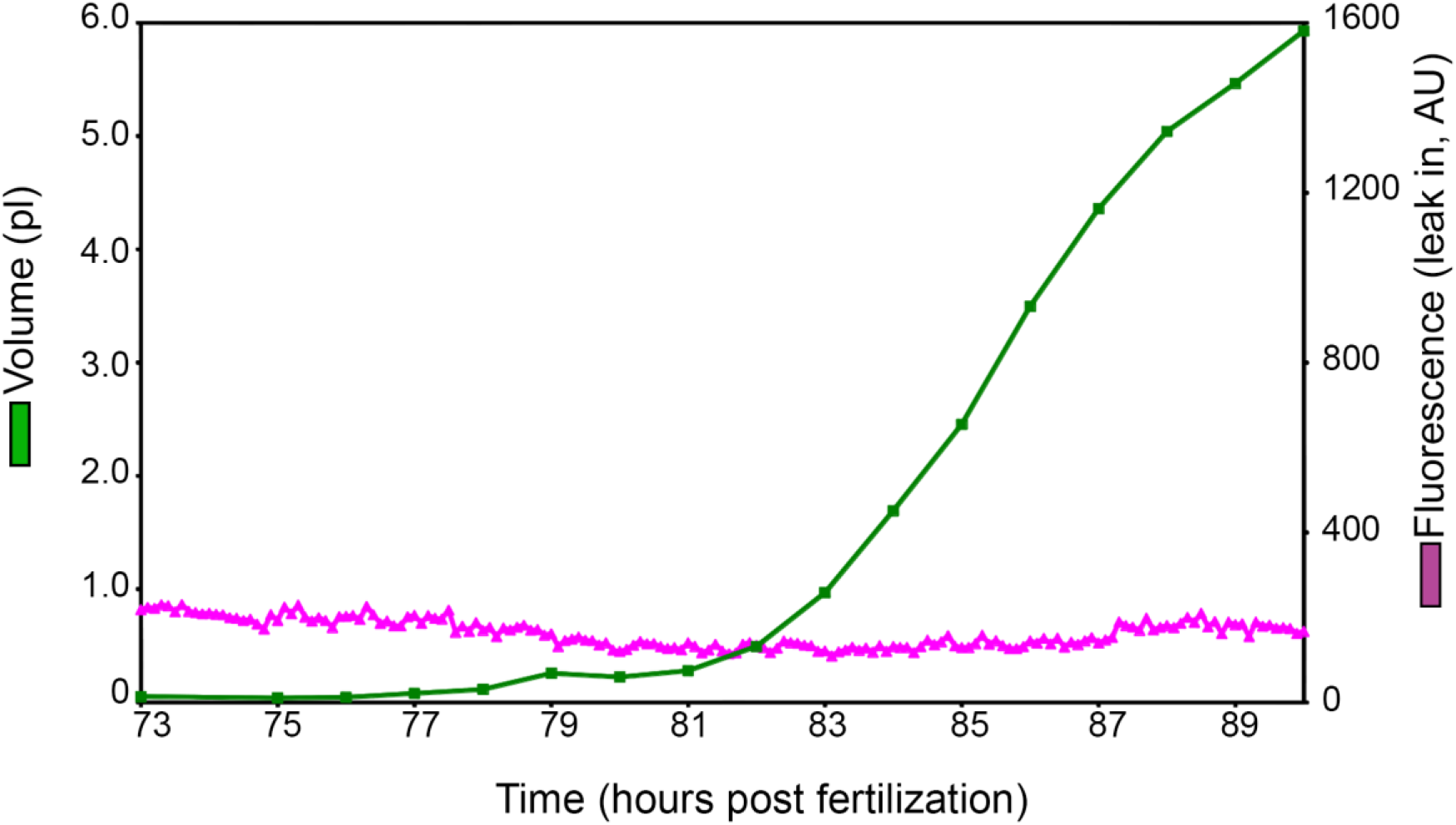
Inflation of additional mutant ES. Quantification of segmented ES volumes (primary axis, green) and leak in fluorescence (secondary axis, magenta) from an additional time-lapse of an *lmx1bb^jj410/jj410^* mutant (see Video 6).

## Videos

**Video 1. Early ES development.** Video begins with schematic of experimental set-up and context of the presented field of view. Then, an annotated time point is presented of the upcoming video. The presented video is of a sagittal slice from a 4D time course of early ES development (white arrow points to ES in introduction). ES morphogenesis begins at 36 hours post fertilization (hpf) as an evagination in the dorsal-anterior-lateral epithelial wall of the otic vesicle. Fluorescence from membrane citrine, shown in grey. Scale bar is 10 μm.

**Video 2. Wild-type ES inflates and deflates.** Video begins with an illustration depicting the context of the presented field of view, which is a sagittal slice encompassing the developing ES from a 4D time course. Then, an annotated time point is presented of the upcoming video, a green arrow points to the ES, a dotted line outlines the ES lumen, the otic vesicle is labeled ventral to the ES, and the perilymph surrounds the ES structure, labeled in magenta. Video of sagittal slice from 4D dataset, quantified in Figure 1F. Fluorescence from membrane citrine, shown in green. Perilymph highlighted with fluorescence from 3 kDa dextran-Texas red, shown in magenta. Scale bar is 10 μm.

**Video 3. Wild-type ES inflates and deflates.** A video of two time courses, sagittal slices from 4D datasets, quantified in Figure 1—figure supplement 1B (left) and Figure 1—figure supplement 1C (right). Fluorescence from membrane citrine, shown in green. Perilymph highlighted with fluorescence from 3 kDa dextran-Texas red, shown in magenta. Scale bars are 10 μm.

**Video 4. Endolymph periodically inflates ES and then leaks into perilymph.** Time course of otic vesicle injected with 3 kDa dextran-Texas red at 55 hpf. Panels are transverse volumes of same time course. Left, labeled endolymph presented in grey. Right, labeled endolymph in magenta, membrane citrine in green. Scale bar is 10 μm.

**Video 5. Mutant ES inflates but does not deflate.** Video of sagittal slice from 4D dataset of lmx1bb^jj410/jj410^ mutant-quantified in Figure 3D. Fluorescence from membrane citrine shown in green. Perilymph highlighted with fluorescence from 3 kDa dextran-Texas red, shown in magenta. Scale bar is 10 μm.

**Video 6. Mutant ES inflates but does not deflate.** Video of sagittal slice from 4D dataset of lmx1bb^jj410/jj410^ mutant-quantified in Figure 3—figure supplement 1. Fluorescence from membrane citrine shown in green. Perilymph highlighted with fluorescence from 3 kDa dextran-Texas red, shown in magenta. Scale bar is 10 μm.

**Video 7. Serial-section electron micrographs of wild-type ES at 5.5 dpf.** Sections are 60 nm thick and color overlays highlight cells with lamellar junctions or basal lamellae. Scale bar is 500 nm.

**Video 8. Slab view of ES time course acquired with lattice light-sheet microscopy with adaptive optics.** 15 sequential slices (300 nm slice spacing) were combined as a maximum intensity projections (MIP) to make a 4.5 μm slab. 7 sequential 4.5 μm slabs were tiled to consolidate the presentation of a complete 3D time course. The membrane citrine signal is green and the 3 kDa dextran-Texas red perilymph highlighter is magenta. Lower right panel is an annotated reference, with the basal surface of the ES labeled and outlined with a dotted yellow line, the apical interface enclosing endolymph outlined with a dotted blue line, the endolymph within the ES lumen indicated with a blue arrow, exposed basal lamellae indicated with a green arrow, and the perilymph labeled with magenta text. Scale bar is 10 μm.

**Video 9. 3D rendering of tracked and segmented cells and ES lumen.** An anterior view on the left and dorsal view on the right. Segmented ES lumen is colored magenta. All other objects are ES cells. Labeled cubes indicate body axes. Same time course as Video 9.

**Video 10. 3D rendering of membrane citrine signal.** The video begins with an annotated time point from the 3D rendering of signal from an AO-LLSM time course. A yellow dotted line highlights the rendered ES, green arrows point to basal lamellae, and the surrounding space is labeled as perilymph with magenta text. Dorsal view of ES, membrane citrine signal rendered in 3D. Scale bar is 10 μm.

**Video 11. Representative heat shock dnRac1 time course.** Embryos were heat shocked at 55 hpf. At 57 hpf, α-bungarotoxin protein and 3 kDa dextran-Texas red were injected into the hearts. Time course began at 58 hpf, membrane citrine is green, perilymph is magenta. Scale bar is 10 μm.

